# Two pore domain THIK2 channel is involved in acute and chronic pain signal regulation

**DOI:** 10.1101/2025.08.08.669258

**Authors:** Nicolas Gilbert, Franck Chatelain, Solène Gibaud, Thomas Lorivel, Ying-Ling Shen, Marie-Emmanuelle Kerros, Sylvain Feliciangeli, Frederic Fiore, Chih-Cheng Chen, Florian Lesage, Delphine Bichet

**Affiliations:** Université Côte d’Azur, CNRS, INSERM, Institut de Pharmacologie Moléculaire et Cellulaire (IPMC), 06560 Valbonne, France; LabEx Ion Channel Science and Therapeutics (ICST), 06560 Valbonne, France; Institute of Biomedical Sciences, Academia Sinica, 128 Academia Road, Section 2, Taipei 115, Taiwan; Taiwan Mouse Clinic - National Comprehensive Mouse Phenotyping and Drug Testing Center, Academia Sinica, Taipei 115, Taiwan; Centre d’Immunophénomique (CIPHE), Aix Marseille Université, INSERM, CNRS, CELPHEDIA, PHENOMIN, 13288 Marseille, France

**Keywords:** K_2P_ channel, Excitability, Peripheral sensory fibers, Knock-out mice, Inflammatory pain

## Abstract

Two-pore domain potassium channels (K_2P_) regulate neuronal excitability by acting as hyperpolarizing leak channels. Among them, THIK2 remain poorly characterized. Although no study has yet clearly linked them to excitability or pain, their selective expression in nociceptive neurons of the Dorsal Root Ganglia (DRG) suggests a role in nociception and pain regulation. This project investigates THIK2 channels in pain pathophysiology through molecular, electrophysiological, and behavioral approaches. We mapped THIK expression patterns and in THIK2 knock-out mice, we examined DRG neuron excitability and pain sensitivity. Results reveal thermal hypersensitivity under both naive and inflammatory conditions, indicating that THIK2 normally limits neuronal hyperexcitability. These findings position THIK2 as a potential therapeutic target in chronic inflammatory pain, with peripheral inhibition potentially offering analgesia without central opioid side effects.

## 1. Introduction

Chronic pain has reached epidemic levels, and its treatment continues to rely heavily on non-steroidal anti-inflammatory drugs (NSAIDs) and opioids. However, these drugs are associated with numerous side effects, including gastrointestinal complications, nephrotoxicity, cardiotoxicity, constipation, and respiratory depression, as well as the risks of tolerance and dependence (1–3). Identifying new pharmacological targets is therefore critical to developing safer and more effective analgesics that minimize adverse effects. Ion channels have emerged as a major class of drug targets for modulating pain due to their presence in sensory neurons within the peripheral and central nervous systems and their pivotal role in regulating cell excitability (4).

In regard to nociception, ion channels are categorized into pro-nociceptive and anti-nociceptive. Pro-nociceptive channels include sodium and calcium channels, transient receptor potential (TRP) channels, acid-sensing ion channels (ASIC), piezo channels, and purinergic P2X receptors (5). Those depolarizing channels initiate and favor the propagation of action potentials in sensory neurons and have been extensively studied for their therapeutic potential in pain management (6).

Several ion channels are now recognized as key mediators of pain transduction and modulation, including TRP, P2X, ASIC, and TACAN channels. Among these, TRP channels play a major role in chemical and temperature detection and thermal allodynia (7–12). ASICs channels have also been identified as mediators of pain induced by tissue acidosis, and their blockade reduces inflammatory hyperalgesia (13–18). P2X receptors are involved in inflammatory pain (19–22). TACAN, identified by Dr. Reza Sharif-Naeini’s team, is a novel mechanically activated channel involved in transducing painful mechanical stimuli, and its inhibition attenuates mechanical nociception (23). Together, these studies highlight the diversity of channels involved in the various components of pain and pave the way for analgesic strategies that specifically target each modality.

Recently, the peripheral sodium channel blocker suzetrigine has demonstrated efficacy in a phase III trial for moderate-to-severe acute pain and was approved as a first-in-class non-opioid analgesic (24). In general, modulators of ion channels are developed for conditions, such as arrhythmias or epilepsy, and their narrow therapeutic margins pose significant challenges for their use against pain. These findings highlight the need to continue exploring ion channels as therapeutic targets for pain management, whether as standalone or combination therapies. Potassium channels belong to the anti-nociceptive group, whose activation inhibit action potentials, decreasing the intensity of painful messages. They represent another promising target for pain relief through the development of channel activators. However, it remains essential to target channels with highly specific expression limited to nociception-related structures to maximize efficacy and minimize off-target effects.

K_2P_ channels represent attractive target due to their unique role in membrane hyperpolarization and their specific expression in pain signaling pathway (25). They generate background potassium currents responsible for the negative cell resting membrane potential (RMP) exerting a brake over cellular excitability. In sensory neurons of the PNS, this brake is essential to counteract action potential lowering or preventing nociceptive signal to be transmitted (26). K_2P_ channels constitute a subfamily of potassium channels and their subunits are encoded by 15 genes (KCNK) divided into 6 groups: TREK/TRAAK, TWIK, TASK, TALK, THIK and TRESK. Functional K_2P_ are made of 2 subunits which can form homomeric and heteromeric channels with different properties (27, 28). Although K_2P_ channels are expressed in a wide variety of cell structures and types, some have been particularly described in peripheral sensory neurons sensitive to mechanical, thermal and potentially injurious stimuli. The peripheral free nerve endings arborize in the skin, muscles, vascular system, viscera, etc., their cell bodies sit in ganglia such as the trigeminal ganglia (TG) or the dorsal root ganglia (DRG), while they make synapses on spinal cord neurons (29). In mammals, peripheral lesions activate nociceptors, generating pain signals transmitted along nerve fibers to TG or DRG, where they undergo initial processing and integration called peripheral sensitization (30). This primary sensory information is subsequently conveyed by neurons to higher centers, including the spinal cord, specific brainstem nuclei, and the cerebral cortex. Within the CNS, these signals are further integrated and amplified by repetitive stimulation, inflammation or injury contributing to the phenomenon known as central sensitization and in the manifestation of chronic pain.

There is already evidence supporting the involvement of certain K_2P_ channels in nociception and pathological pain (25). Several K_2P_ channels have been identified in TG and DRG, where they modulate pain signaling. TREK1 channels for example colocalize with TRPV1 and substance P, and both TREK1 and TREK2 are associated with IB4+ fibers (31, 32). TRAAK appears to be expressed indifferently in all sensory fibers (31, 33). Their presence in these neurons has been linked to the modulation of their excitability and their pattern of spike firing (32, 34, 35). Behavioral studies from TREK/TRAAK KO mice have also demonstrated altered sensitivity to mechanical and/or thermal stimuli in acute and chronic pain models (34). More interestingly, agonists targeting TREK1 channels have demonstrated analgesic effects but with potential side effects probably due to its wide expression in other cells and particularly in cardiomyocytes and neurons form central nervous system (36–38). Although existing potassium channel activators demonstrate potential analgesic effects, they were originally developed for other therapeutic indications, and their use in pain treatment is constrained by the risk of adverse side effects. In addition to TREK/TRAAK, TRESK channels have also been widely expressed in sensory neurons from both DRG and TG (31, 39, 40). Behavioral studies using KO mice or viral treatments to inhibit TRESK in sensory neurons have not shown a significant impact on general sensitivity to mechanical and thermal stimuli, except when the stimuli were applied to the face under acute or pain-induced conditions (such as capsaicin or acetone cheek injections) (41, 42). Aside from TREK/TRAAK and TRESK channels (34, 43–49), there is limited evidence supporting the involvement of other K_2P_ channels, such as those from the TASK and TWIK groups, in pain modulation (50, 11, 51–53). The main limitation to further exploration in this area is the lack of selective modulators for these channels, despite research is focusing on the development of modulators of K_2P_ channels (54).

We developed THIK2 knockout (THIK2^−/−^) mice by genetically deleting the THIK2 channel to investigate its role in acute and chronic pain. Characterization of this new model revealed that THIK2 is predominantly expressed in non-peptidergic C-type DRG neurons, and its absence affects the excitability of these specific neurons. Using in vivo behavioral tests, we compared mechanical and thermal sensitivities under both naïve and chronic conditions. THIK2 KO mice demonstrated increased sensitivity to thermal stimuli in the naïve state and exhibited both thermal and mechanical hyperalgesia under inflammatory conditions. Furthermore, these mice displayed elevated pain responses in the spontaneous formalin pain test and showed delayed recovery from inflammation. These findings position THIK2 as a promising new target for pain treatment.

## 2. Material and Methods

### Mouse strain

All protocols are in agreement with national and European Union recommendations 2010/63/EU with approval of the Animal Care Committee of Nice for animal experimentation and have been approved by “MINISTERE DE L’ENSEIGNEMENT SUPERIEUR ET DE LA RECHERCHE” under the reference number: APAFIS #37438-202205131616821 v7 and APAFIS #44595-2023073117435391 v6. Experiments were performed on wild-type (WT) C57BL/6J mice (Janvier labs, France) and THIK2 knockout mice (THIK2^−/−^) from C57BL/6J mice. THIK2^−/−^βGal^+/+^ mice were generated in Centre d’Immunophénomique (CIPHE) in Marseille, by knocking down Kcnk12. the gene encoding for THIK2 and by replacing this gene by LacZ, encoding for β-galactosidase. We observed no compensatory upregulation of genes for other neuronal K_2P_ channels in DRGs of THIK2^−/−^βGal^+/+^ mice, particularly THIK1, that is the closest in structure to THIK2 (Supplementary Figure S1A). THIK2^−/−^ mice also express essentially unchanged levels of TRPs and ASICs channels (Supplementary Figure S1A) except a decrease for ASIC3.

### Animals Housing

Males and Females C57Bl/6j mice were housed in groups of 4 or 5. under standard conditions (20-22 °C, 40% humidity), with 12h light/dark cycle (lights off at 8 P.M.), with free access to food and water. The mice used were all C57Bl/6J mice, separated into 3 groups, THIK2^−/−^, WT mice purchased from Janvier labs and WT littermate mice. They were all tested between 8 and 12 weeks old. Efforts were made to minimize the stress and potential suffering of mice used in this study.

### Nociceptive behavioral assays

All behavioral experiments were performed on 20–25g C57Bl/6J mice between 8 and 12 weeks old. Animals were acclimated for 45min to their testing environment prior to all experiments that are done. Experiments were made at room temperature (20-22 °C) in a quiet room and were performed blind to the genotype. The number of tested animals is indicated in the figure legends section. Von Frey and Hargreaves apparatus were from UgoBasile, Italy.

### Formalin spontaneous pain test

Mice were placed individually into mirror chambers 45 min before injection to allow them to acclimatize to their surroundings. Following intra-plantar injection of 10 μ l of a 2% formalin in NaCl 0.9% solution with a 31-gauge needle into the left hind paw, four nociceptive behaviors were quantified: time spent shaking, lifting, licking or biting the injected paw was monitored for 45 min and analyzed for 5 min intervals. This test can be separated in two phases: an initial acute phase I (0-10 min) followed by a relatively short quiescent period, and then started a prolonged tonic response (phase II from 10-45 min) (55).

### Post formalin recovery test

A Noldus® Phenotyper was used to analyze rearing and resting behavior. The phenotyper is equipped with a camera and the videos were analyzed using Noldus® Ethovision software. The animals were placed for 1 hour, without food or water, in a 30×30cm box with clean bedding. This test was carried out a first time 10 days before the formalin test (naive condition), and a second time 1 hour after formalin 2% injection into the hind paw (post-formalin condition).

### Mechanical sensitivity Von Frey test

Mice were placed individually into chambers with a wire mesh grid floor 45 min before testing. Mechanical allodynia was assessed using calibrated Von Frey filaments ranging from 0.16 to 2g and dynamic Von Frey apparatus with ramped increasing force (0 to 7.5g) from Ugo Basile, Italy. For the manual test, the filaments, tested in order of increasing stiffness, were applied 5 times, at more than one-minute interval, perpendicularly to the plantar surface of the hind paw and pressed until they bent. Following the “ascending stimulus” method, the threshold was determined following 4 consecutive positive responses to the same filament (56).

### Radiant heat Hargreaves test

Mice were placed individually into chambers with a plexiglass floor 45 min before testing. Thermal radiant heat was assessed using the Ugo Basile Hargreaves device calibrated at 50% and 60% of power. 50% power (260mW/cm^2^) was used to produce withdrawal latencies of 10-12s in naïve animals. The threshold was determined by averaging 4-8 values for each hind paw of each animal.

### Complete Freund’s Adjuvant (CFA)-induced pain model

We made an intra-plantar injection of 15 μl of a 1:1 CFA (ThermoScientific) in saline emulsion with a 26-gauge needle. We then measured the thermal and mechanical thresholds as described above every 2 or 4 days until D+28 post-CFA injection.

### X-Gal staining

To obtain adult tissues, five male THIK2^−/−^βGal^+/+^ mice were anesthetized with isoflurane (Vetflurane®) and decapitated. Then L1 to L6 DRGs were dissected and were placed in a solution containing X-Gal, which reacts with the enzyme β-galactosidase. After one day, images of whole DRG were taken using macroscope. DRG were finally placed in cryoprotective solution (OCT) before being frozen and stored at −80°C. DRG were sectioned at 10 μm using a standard cryostat (Leica). Brightfield Images of the slices were taken on a 20X air objective on histology DMD108 Leica microscope. Lumbar DRG from THIK2^−/−^βGal^+/+^ mice were counted in five independent animals. All cell counts were conducted by an individual on OMERO software. Statistical significance was set to p < 0.05 and assessed using unpaired T-test.

### Fluorescent situ hybridization (FISH)

In situ hybridization and immunofluorescence were carried out following RNAscope Multiplex V2 protocol for Fresh Frozen tissues form Bio-techne®. To obtain adult tissues, mice were anesthetized with isoflurane (Vetflurane®) and decapitated. Then, L1 to L6 DRGs were dissected, immediately placed in cryoprotective solution (OCT) before being frozen and stored at −80 °C. DRG were sectioned at 10 μm (DRG section) using a Leica CM3050S cryostat. In situ hybridization was performed with the Bio-techne® RNAscope Multiplex V2 in accordance to the manufacturer’s protocol. Specific probes were used to detect Kcnk12 (THIK2), Kcnk13 (THIK1), P2rx3 (Purinergic receptor X 3), Cgrp (Calcitonin Gene-Related Peptide), Th (Tyrosine Hydroxylase) and Nf200 (NeuroFilament 200) mRNAs. Hybridization with a probe against the Bacillus subtilis dihydrodipicolinate reductase (dapB) gene (Ref 320871) was used as a negative control and a mix of PPIB, UBC and Pol2RA (Ref 320881) was used as a positive control. Probes were hybridized 2h at 40 °C. Final detection was achieved using TSA Vivid kit 520. 570 and 650 (TOCRIS). Z-stack Images of these fluorophores on a laser scanning confocal microscope (TCS SP8 Leica microsystem) through a 63X objective. Quantifications were done on maximum Z-stack projections using QuPath v0.5.0 software (57), area was automatically determined by the software after identification of the labelled neurons. Then, the number of dots per cell was detected after applying the same detection threshold to all images. For each genotype, lumbar DRG were counted in three independent animals. Statistical significance was set to p < 0.05 and assessed using Two-way ANOVA analysis followed by post-hoc Tukey test.

### DRG culture and patch clamp recordings

DRG neurons were obtained from 8 to 12-week-old wild type (C57Bl/6J) and THIK2-/- mice. After isoflurane anesthesia, animals were killed by decapitation and lumbar DRG were retrieved. Collagenase enzymatic digestion and slow mechanical dissociation were made. DRG neurons were plated in 35 mm petri dishes previously coated with poly-(L)-lysine and cultivated in complete Neurobasal. Patch-clamp experiments were performed between 1 to 3 days after plating. IB4 (1:2000) conjugated with AlexaFluorR 488 is applied 4 hours before patch-clamp recordings. Non peptidergic C-fiber neurons were determined by Isolectin B4 staining. All recordings were made on the same electrophysiological recording station, at room temperature (20-22 °C). External solution contained (in mM): 140 NaCl, 4 KCl, 1 CaCl2. 1 MgCl2. 10 HEPES, (pH 7.4 adjusted with NaOH). Patch pipettes had a resistance of 3.5–5.5 MΩ and were filled with (mM) 155 KCl, 4 MgATP, 5 EGTA and 10 HEPES (pH 7.2 with KOH). Voltage was observed by using 700B Multiclamp (Axon Instruments), low-pass filtered Bessel at 1 kHz, and digitized at 10 kHz by using a Digidata-1550B (Axon Instrument). Clampex was used for current visualization and stimulation. Current-clamp: Membrane potential was recorded under control bath solution perfusion and under bath solution perfusion. Rheobase was obtained after successive application of 20 pA until an action potential was obtained. The protocol then consisted of applying 10 sweeps, each sweep consisting of stimulation with a current equal to 3x the rheobase for 5 ms to trigger an action potential, then after 500 ms of rest, we again applied a current equal to 3x the rheobase for 800 ms to trigger trains of action potentials, followed by 1 s of rest. Data were obtained and analyzed with Clampfit software. Statistical significance was set to p < 0.05 and assessed using unpaired T-test or Mann-Whitney test.

### Statistical analysis

Statistical significance was set to p < 0.05 and assessed using unpaired T-test analysis (Hargreaves, dynamic Von Frey), one-way ANOVA analysis followed by post-hoc Dunett test (for area under curve of the formalin test), two-way ANOVA followed by post-hoc Dunett and Bonferroni test (for thermal and mechanical allodynia with CFA), two-way ANOVA followed by post-hoc Tukey test (for formalin test, manual Von Frey and post formalin recovery) using GraphPad Prism 10.4.1 software. All error bars represent standard deviation (SD) or standard error of the mean (SEM).

## 3. Results

### THIK1 and THIK2 channel expression in DRG

Transcriptomic analyses of mouse trigeminal ganglia (TG) and dorsal root ganglia (DRG) have identified THIK2 as the most abundantly and specifically expressed K2P channel in human (58, 59). However, its functional role in these neurons, as well as its broader physiological significance, remains unclear. In humans, THIK2 is also the most highly expressed K_2P_ in DRG and TG neurons, surpassing even TREK1 and TRESK. In fact, THIK2 ranks among the ten most expressed ion channel transcripts in peripheral sensory neurons (58). To study the expression of THIK2 in the peripheral sensory system and investigate its role in the regulation of sensory signals, we created a new reporter mouse carrying the gene for β-galactosidase in place of *Kcnk12* gene coding THIK2 subunit. We collected adult mice ganglions from hetero and homozygous mice and used WT as negative control. The expression of β-gal was investigated by X-gal staining on whole lumbar and thoracic DRGs from homozygous THIK2^−/−^βGal^+/+^ mice (Fig. 1C) whereas WT littermate served as negative control (Supp. Fig. 2). We observed clear labeling of all ganglia in βGal+ mice, appearing as dot-like patterns at the ganglion level, resembling the labeling of sensory neuron cell bodies. A more detailed analysis by sectioning the DRG (10μm) showed that X-Gal staining was present in only some structures and not all cells (Fig. 1C). We quantified X-Gal positive and negative cells from 1 DRG per mouse from 5 mice and analyzed staining relatively to the cell body area (in μm^2^). Size distribution analysis showed presence of X-Gal staining with a higher proportion in small and medium-sized neuron compared to X-Gal negative neurons. The averaged sized of X-Gal positive neurons being 549±289 μm^2^ compared to 897±489 μm^2^ for X-Gal negative (mean ± SD, Fig. 1C). Although there are more than two subtypes of peripheral sensory neurons, the basic classification divides them into fast-conducting A-fibers and slow-conducting C-fibers. The size difference between A-fibers and C-fibers primarily relates to their diameter and conduction velocity, which are influenced by their myelination and function. A-fibers diameter ise typically larger than 650 μm^2^ in cell body area whereas C-fibers diameter is typically smaller than 650 μm^2^. Using X-Gal staining on DRGs from THIK2^−/−^βGal^+/+^ mice, we demonstrated that THIK2 is mostly expressed in small size neurons (<650 μm^2^) corresponding to C-fibers (Fig. 1C).

**Fig. 1.**
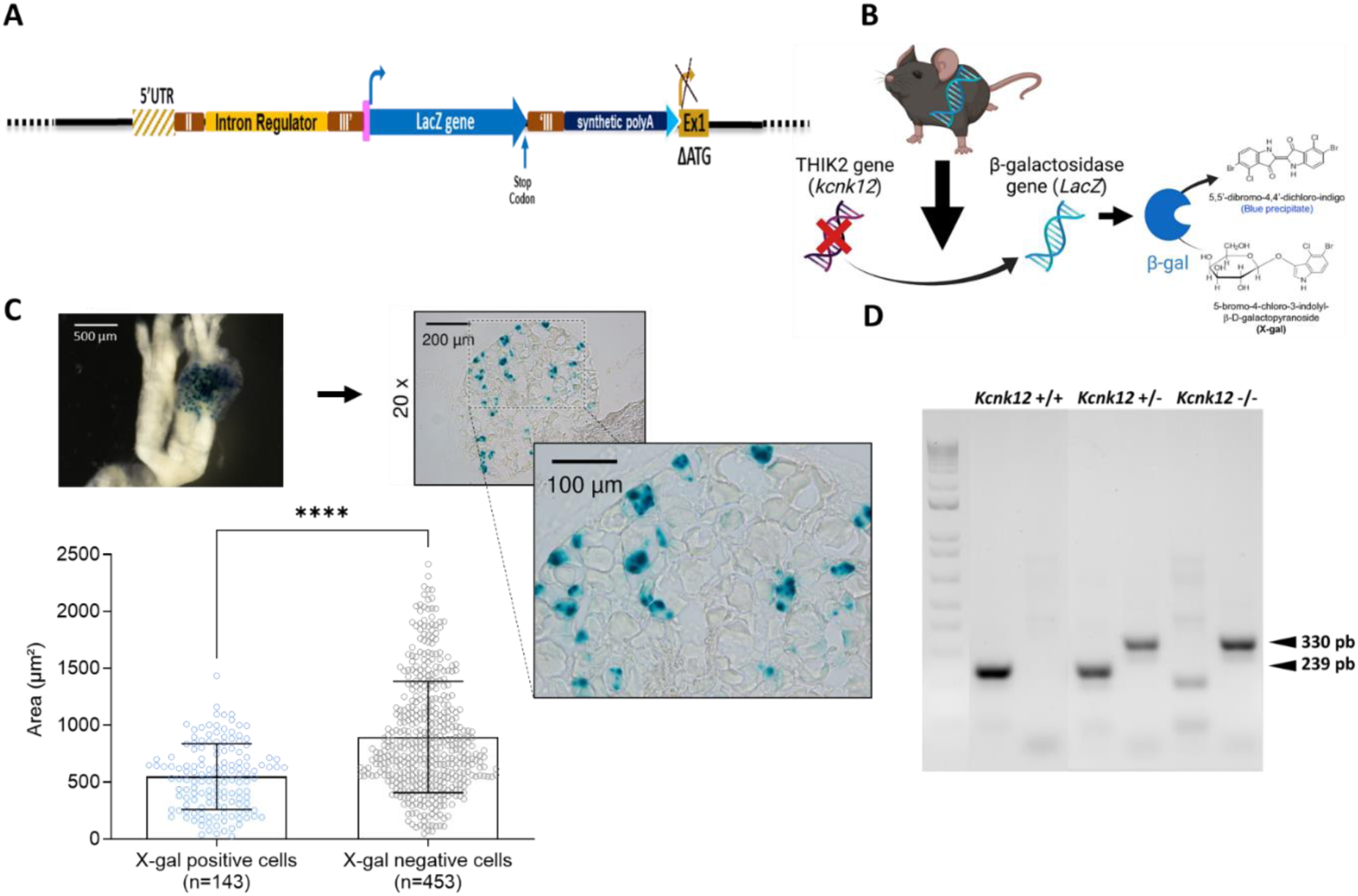
X-Gal staining in THIK2^−/−^Gal^+/+^ mouse lumbar DRG neurons. **A.** The *Kcnk12*^−/−^ was made by introducing the LacZ gene in the first exon after the first ATG. **B.** The LacZ gene codes for β-galactosidase (β-gal) in *Knck12^−/−^* mice which turns blue in response to the X-Gal compound**. C.** Images of β-gal labeled whole lumbar DRG from 5 different THIK2^−/−^Gal^+/+^ homozygous male mice. DRG cryostat sections are 10 μm thick and magnifications used are 20X and 40X magnification. Quantification of β-gal positives (n=143) and negatives (n=453) cells. Mean ± SD, unpaired T-test, ****p<0.0001. **D.** Genotyping of *Kcnk12*^+/+^ (239 pb), *Kcnk12*^+/-^ and *Kcnk12*^−/−^ (330 pb) mice.

DRG neurons can also be molecularly characterized by their mRNA expression profiles, with specific markers commonly used alongside size to identify nociceptor subpopulations. Traditionally, these neurons are grouped into four main categories: CGRP-positive peptidergic nociceptors, P2XR3-positive (also IB4-positive) non-peptidergic nociceptors, TH-labeled C-low threshold mechanoreceptors (C-LTMR) and large myelinated neurons expressing neurofilament-200 (NF200) (60). To further distinguish between fiber types (A or C) and specifically identify peptidergic versus non-peptidergic C-fibers, we employed RNAscope (fluorescent in situ hybridization, FISH) on WT DRGs. This approach allowed us to examine THIK2 channels in conjunction with markers such as NF200, P2RX3, CGRP and TH. Our findings revealed that THIK2 (in red) is predominantly expressed in P2RX3-positive neurons (in green), corresponding to non-peptidergic (NP) nociceptors (Fig. 2A). These results align closely with RNA-seq data in mice, which demonstrate that THIK2 transcripts are the most abundant among K_2P_ transcripts in non-peptidergic C-fibers (59). We quantified THIK2 expression in the four labeled neuronal populations by counting the number of dots per cell corresponding to THIK2 labeling (Fig. 2B). This analysis revealed that P2RX3 neurons (89.72±7.34 dots/cell) exhibited THIK2 levels approximately 5 times higher than CGRP neurons (17.16±5.12 dots/cell), with minimal to no detectable THIK2 labeling in TH (4.29±1.17 dots/cell) and NF200 (3.03±0.82 dots/cell) neurons.

**Fig. 2.**
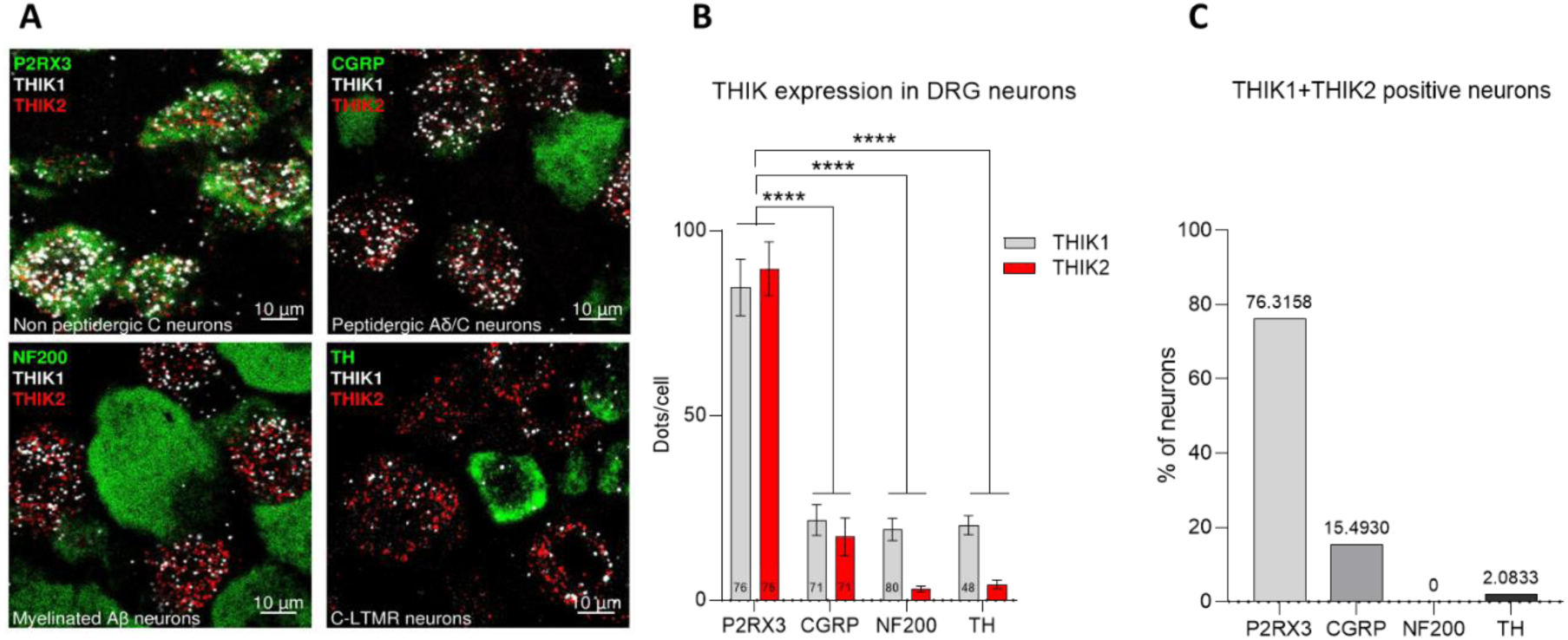
THIK mRNA expression in WT mouse DRG neurons. **A.** Confocal images of mouse lumbar DRG sections co-labeled with mRNA probes for THIK2 (red), THIK1 (white) and neuronal markers for non peptidergic neurons (P2RX3), peptidergic neurons (CGRP), myelinated large neurons (NF200), and C-Low Threshold Mechanoreceptor neurons, C-LTMR (TH). **B.** Quantification of THIK2 and THIK1 mRNAs dots in P2RX3 (n=76), CGRP (n=71), NF200 (n=80) and TH (n=48) positives DRG neurons from 3 independent male mice. Mean ± SEM, Two-way ANOVA with Tukey’s multiple comparisons test, ****p<0.0001. **C.** Neurons positivity (>20dots/cell) percentage for different markers.

This finding is particularly noteworthy due to the specificity of this channel for non-peptidergic C neurons. We previously demonstrated that THIK1 and THIK2 can form functional heterodimers (61). To further investigate their relationship, we used a THIK1 RNA probe (in white) to assess THIK1 expression relative to THIK2. Interestingly, our RNAscope experiments revealed that both THIK1 and THIK2 are co-expressed in P2RX3-positive neurons, accounting for more than 76% of the cells (Fig. 2C). Additionally, THIK1 displays an expression pattern similar to THIK2, indicating potential homo- and/or heterodimer formation between the two channels in these neurons (Fig. 2A-B). We also explored whether the absence of THIK2 alters the expression of THIK1 or other ion channels. Using RNAscope, we re-evaluated THIK1 RNA expression in DRG sections from THIK2 KO mice. Our analysis showed no increase in expression patterns; instead, we observed a statistically significant decrease in THIK1 levels in P2RX3, CGRP and NF200 neurons (Supplementary Figure S1B). This finding was corroborated by Biomark® qPCR and extended to include other ion channels from the K_2P_ family, as well as channels implicated in pain regulation, such as ASIC and TRP channels, only a decrease ASIC3 is remarkable (Supplementary Figure S1A). The minimal changes observed in the expression of other genes following THIK2 deletion underscore the specificity of THIK2’s function. With the possible exception of a compensatory interaction with THIK1, these findings support the robustness of the animal model for investigating THIK2’s role, free from significant confounding effects due to broader compensatory mechanisms.

### Increased excitability of non peptidergic DRG neurons from THIK2 KO mouse

If the THIK2 channel influences the excitability of sensory neurons, we would expect to observe variations in neuronal activity depending on its presence or absence. To explore this, we performed electrophysiological studies on DRG neurons cultured from our THIK2^−/−^βGal^+/+^ (THIK2 KO) mouse model. These experiments were conducted on 2-3-days neuronal cultures from male, adhering to established protocols. Using RNAscope, we identified that THIK2 is predominantly expressed in non-peptidergic C neurons. Based on this, we chose to record from small to medium-sized neurons, likely originating from C-type fibers, as well as IB4+ neurons to specifically target non-peptidergic C-nociceptors. The membrane capacitance, directly proportional to the cell surface area, was measured using the voltage-clamp technique. The results indicated no significant difference between WT and THIK2^−/−^ neurons, suggesting comparable cell sizes (WT: 39.89 ± 2.86 pF, n = 26; THIK2^−/−^: 41.61 ± 2.16 pF, n = 22; p = 0.648) (Fig. 3E). Switching to current-clamp configuration, we assessed the resting membrane potential (RMP), which was on average, slightly more depolarized in THIK2^−/−^ neurons (−48.41 ± 1.97 mV, n = 22) compared to WT (−50.01 ± 1.62 mV, n = 26), though the difference was not statistically significant (p = 0.509) (Fig.3F). Input resistance was measured as 7.07 ± 0.47 MΩ for THIK2^−/−^ and 6.95 ± 0.59 MΩ for WT neurons (Fig. 3G). Neither THIK2^−/−^ nor WT DRG neurons exhibited spontaneous action potential (AP) firing at RMP. To investigate membrane excitability, we applied incremental depolarizing current steps (1500 ms, 20 pA increments) starting from 0 pA mV to evoke a single AP and determined the rheobase, the smallest current required to elicit an AP. The rheobase was slightly higher in THIK2^−/−^ neurons (77.73 ± 10.73 pA, n = 22) compared to WT (70 ± 9.10 pA, n = 26) but not significant (p=0.476) (Fig. 3H). The AP threshold was also slightly different between the two groups (THIK2^−/−^: −18.5 ± 1.10 mV; WT: −15.5 ± 1.21 mV) but not significant (p=0.077) (Fig. 3I). However, during prolonged stimulation (800 ms), which allowed for AP train generation, THIK2^−/−^ neurons exhibited a significant (p=0.0467) higher number of APs (6.45 ± 0.81) compared to WT neurons (4.81 ± 0.96) (Fig. 3C, D and J). These results demonstrate that THIK2 significantly contributes to small IB4-positive DRG neurons excitability. Notably, 95.45% of THIK2^−/−^ neurons displayed at least two AP responses, whereas only 57.69% of WT neurons did so. AP parameters, assessed using stimulation at three times the rheobase, including peak potential and AP duration, showed minimal or no differences between WT and THIK2^−/−^ neurons (Fig. 3K and L). These findings suggest that THIK2 current plays a limited role in these aspects of AP generation.

**Fig. 3.**
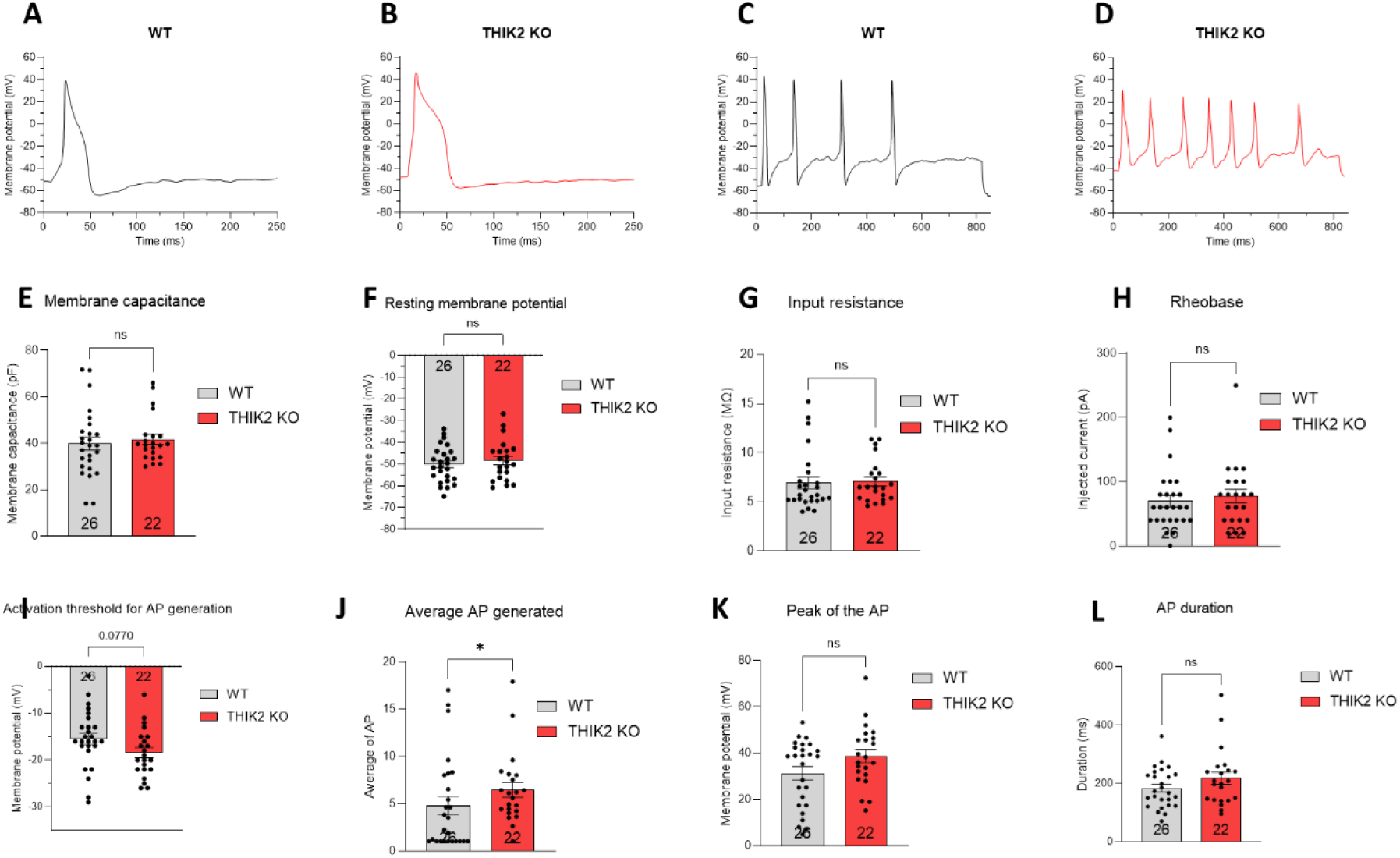
Electrophysiological properties of isolated IB4+ neurons from WT and THIK2^−/−^ Mice. **A-D**. The traces represent action potential generated by depolarizing current injection calculated with 3x rheobase on an isolated neuron, 300pA for the WT and 150pA for the THIK2^−/−^. **E-L**. Membrane capacitance (E), resting membrane potential (F), input resistance (G), rheobase (H), activation threshold (I), average action potentials (J), peak of the action potential (K) as resting amplitude after action potential (L) of non peptidergic IB4+ WT (n=26) or THIK2^−/−^ (n=22) neurons were determined after patch-clamp recordings. Mean ± SEM, Unpaired T-test (E-F-G-H-I-K-L) except Mann-Whitney test for Average AP (J), *p<0.05.

### Acute mechanical and thermal sensitivity of THIK2 KO mice

To investigate whether the hyperexcitability of IB4-positive DRG neurons in THIK2 KO mice translates into a whole-animal behavioral phenotype, we assessed the mechanical and thermal sensitivity of naive mice. Paw withdrawal latencies to mechanical stimuli of increasing weights were measured using dynamic (Fig. 4A) or manual (Fig. 4B) Von Frey tests on separate groups of mice. The manual test involved 48 mice per phenotype, evenly split between males (24) and females (24). The dynamic test was conducted on a second group consisting of 20 WT mice (10 males, 10 females) and 19 THIK2^−/−^ mice (10 males, 9 females). No differences in mechanical sensitivity were observed between THIK2^−/−^ and WT mice, regardless of the test used, with paw withdrawal thresholds recorded as in response to 3.89 ± 0.15 g for WT compared to 4.01 ± 0.16 g for THIK2^−/−^ (Fig. 4A). No gender-related differences were identified (Supplementary Figure S2A-B). The mice were also tested in mechanical induced itch by Von Frey filaments and the results showed no overall difference from WT mice (Supplementary Figure 5A). Only the young (3.5 months old) THIK2^−/−^ mice showed more scratching in response to 0.07 g. The Hargreaves test was performed at 50 % and 60 % of the device’s maximum power on two separate groups of mice. At 50 % power (260 mW/cm^2^), 18 WT mice were tested, including 9 males and 9 females, the results revealed thermal allodynia for THIK2^−/−^ mice (Fig. 4C) but we identified a gender-related difference to thermal sensitivity. The results revealed that at this intensity, thermal hypersensitivity was exclusively observed in females THIK2^−/−^ (Supplementary Figure S2C). When a higher intensity (60 % power) was applied to two larger cohorts 46 WT mice (22 males, 24 females) and 48 THIK2 KO mice (24 males, 24 females), thermal hypersensitivity was observed in both sexes (Supp. Fig. 4). The paw withdrawal thresholds recorded as 50 % intensity was 10.84 ± 0.49 s for WT compared to 9.39 ± 0.48 s for THIK2^−/−^ mice showing nearly 15% of decrease (Fig. 4C). The paw withdrawal thresholds recorded as 60 % intensity was 6.43 ± 0.32 s for WT mice compared to 4.57 ± 0.21 s for THIK2^−/−^ mice, showing almost 30 % of decrease (Fig. 4D). Thermal withdrawal thresholds were systematically lower in THIK2^−/−^ compared with WT control mice (Fig. 4). The data underscore for the first time the importance of THIK2 channels for heat sensitivity and correspond to the first demonstration of the role of a THIK channel in nociception.

**Fig. 4.**
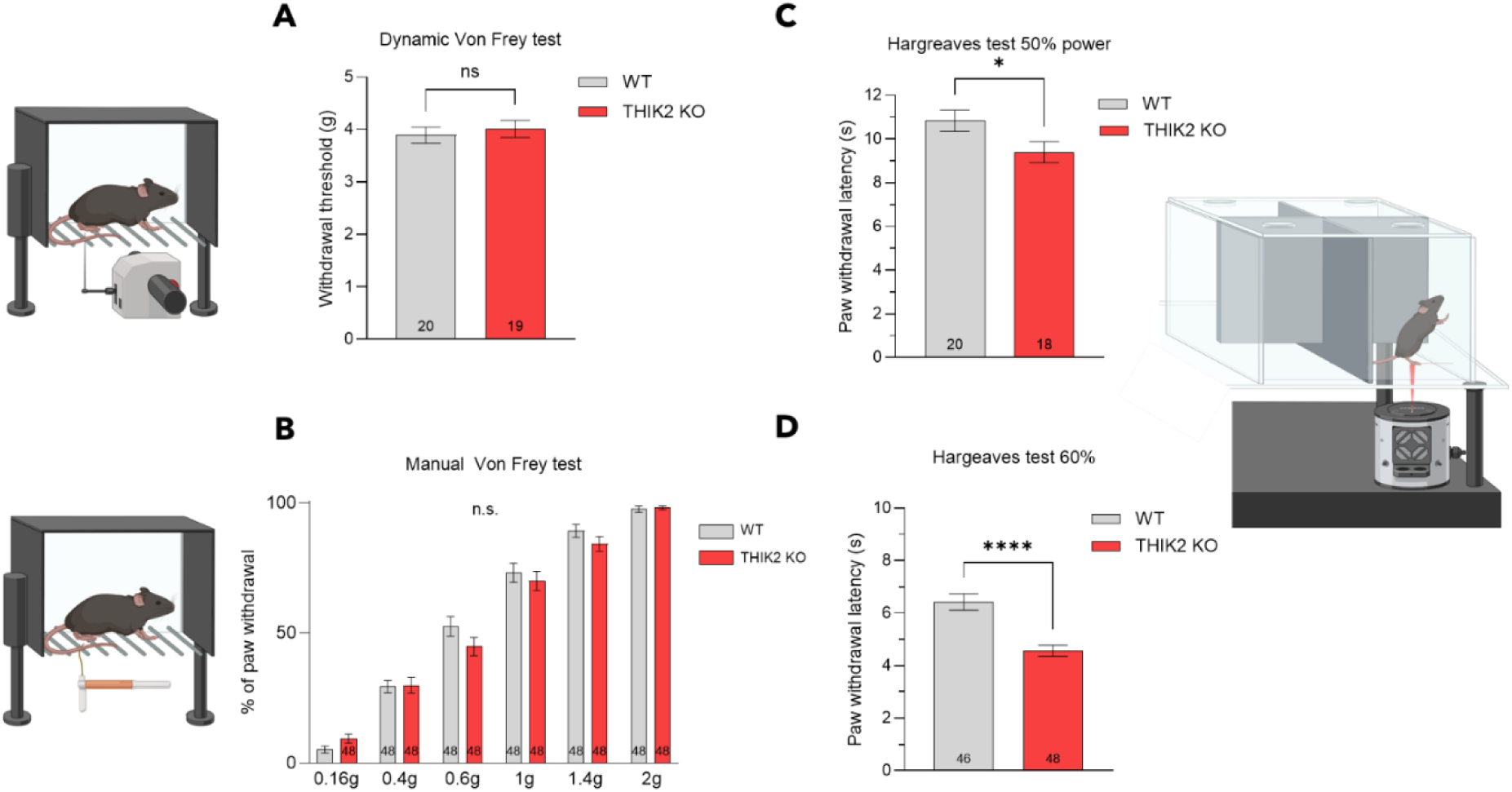
Acute mechanical and thermal sensitivity of WT and THIK2^−/−^ Mice. **A.** Paw withdrawal threshold of THIK2^−/−^ (n=19) compared to WT (n=20) mice in response to mechanical stimuli in dynamic von Frey test (1-7.5g). Mean ± SEM, Unpaired T-test. **B.** Paw withdrawal threshold of THIK2^−/−^ (n=48) compared to WT (n=48) mice in response to mechanical stimuli in manual Von Frey test (0.16-2g). (Mean ± SEM, two-way ANOVA). **C**. Paw withdrawal latency of THIK2^−/−^ (n=20) compared to WT (n=18) mice in response to 50% thermal stimuli intensity in Hargreaves test. Mean ± SEM, Unpaired T-test, *p<0.05. **D.** Paw withdrawal latency of THIK2^−/−^ (n=46) compared to WT (n=48) mice in response to 60% thermal stimuli intensity in Hargreaves test. Mean ± SEM, Unpaired T-test, ****p<0.001

### THIK2 and acute inflammatory pain

Given the expression of the THIK2 channel in polymodal non-peptidergic C-fiber neurons of the DRG, which are involved in inflammatory pain (62), it was necessary to investigate the impact of THIK2 channel deletion on both acute and chronic inflammatory pain. To this end, we conducted different inflammatory pain tests in mice. A spontaneous pain test using 2 % formalin injection in the paw was performed on a batch of 20 WT mice (10 males, 10 females) and 24 THIK2^−/−^ mice (10 males, 14 females), and 13 WT littermates (Supplementary Figure S3). During the acute chemical pain phase (phase 1, <10 min), no significant difference in spontaneous pain behavior was observed between THIK2^−/−^ and WT mice. However, during the acute inflammatory phase (phase 2, 10-45 minutes), THIK2^−/−^ mice exhibited a significant increase in spontaneous pain behavior, such as licking, biting, lifting and shaking their hind paw, with a peak observed at 20 minutes (Fig. 5A), compared to WT mice (and WT littermates, Supplementary Figure S3). The AUC resulting from the increase in the acute inflammatory phase 2 was significantly 30 % higher for THIK2^−/−^ mice, with a value of 2314.03 ± 183.49 for THIK2^−/−^ mice compared to 1643.37 ± 140.87 for WT mice (Fig. 5B). These results confirm the inflammatory hyperalgesia phenotype of THIK2^−/−^ mice. We were also aimed to investigate post-formalin recovery. According to the literature, WT animals typically no longer display spontaneous painful behavior one hour after formalin injection (63). After confirming that THIK2^−/−^ mice also stopped spontaneous painful behavior following the end of the formalin test (Fig. 5C), we sought to determine whether THIK2^−/−^ mice exhibited the same recovery behavior as WT mice. To assess this, animals previously subjected to the formalin test and, 15 minutes after its completion, were placed in a Noldus Phenotyper® observation cage. The experiment included 18 WT mice (10 males, 8 females) and 23 THIK2^−/−^ mice (10 males, 13 females). Following the previous formalin spontaneous pain test (+FORM), THIK2^−/−^ mice exhibit a twofold increase in resting time, spending 952.94 ± 120.96 s at rest compared to 484.52 ± 151.10 s for WT mice (Fig. 5C). Additionally, THIK2^−/−^ mice display a significant reduction in rearing behavior, with only 75.95 ± 6.06 s spent rearing versus 121.10 ± 17.2 9s for WT mice (Fig. 5D). Notably, we observed sex differences, with female THIK2^−/−^ mice showing more pronounced pain behaviors and slower recovery than their male counterparts (Supplementary Figure S3). To evaluate whether this phenotype extended to chemical-induced itch responses, THIK2^−/−^ mice were tested in the cheek skin assay model by intradermal injections of chloroquine or capsaicin. No significant differences were observed in drug-induced wiping (pain-like) or scratching (itch-like) behaviors (Supplementary Figure S5B). Importantly, that difference in pain sensibility is not attributable to increased or altered locomotion. THIK2^−/−^ mice performed normally in both open-field test and elevated plus maze, with no signs of altered locomotion and anxiety levels (Supplementary Figure S6). There was even a slight increase in the distance covered by THIK2^−/−^ mice in the maze. These findings suggest that the observed differences are specific to pain sensitivity rather than general behavioral deficits.

**Fig. 5.**
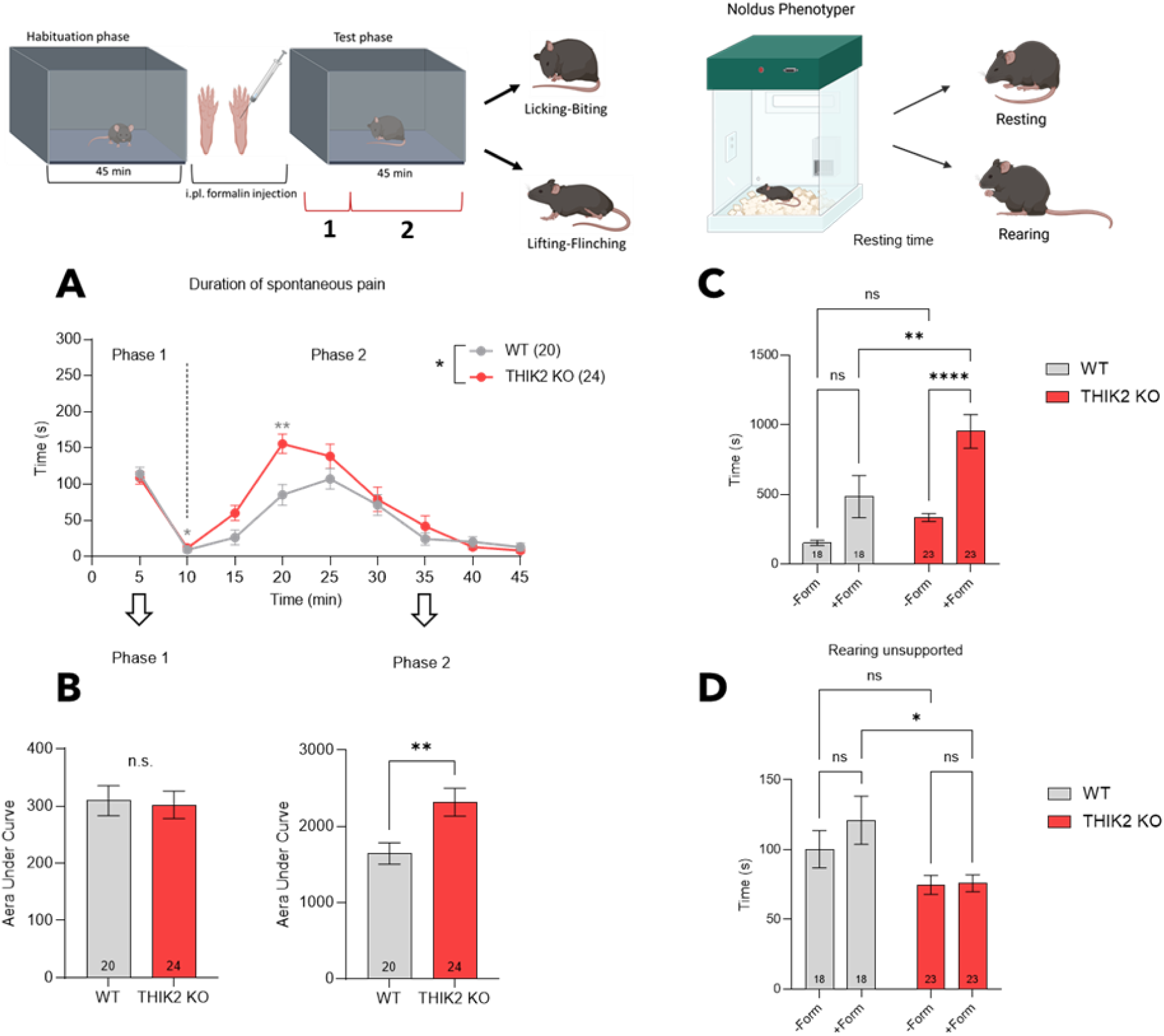
Acute inflammatory spontaneous pain of WT and THIK2^−/−^ Mice. **A.** The pain behaviors measured for WT (n=20) and THIK2^−/−^ (n=24) mice in the formalin spontaneous pain test were licking, biting, shaking and paw lifting. The test lasted 45 minutes after the injection of 15μL of formalin into the left hind paw. Mean ± SEM, two-way ANOVA with with Tukey’s multiple comparisons test, *p<0.05, **p<0.01. **B.** Area under curve form behaviors were separated into two phases: Acute phase between 0-10min and inflammatory phase 10-45min. Mean ± SEM, Unpaired t-test, **p<0.01. **C-D.** Post-formalin recovery was tested on WT (n=18) and THIK2^−/−^ (n=23) mice by placing the animals in a Phenotyper® observation cage 15min after the end of the formalin test, resting/prostration time and time spent rearing were measured. Mean ± SEM, two-way ANOVA with with Tukey’s multiple comparisons test, *p<0.05. **p<0.01. and ****p<0.001.

### THIK2 and chronic inflammatory pain

After demonstrating the involvement of the THIK2 channel in acute inflammation (Fig. 5), we next investigated its role in chronic inflammation pain in THIK2^−/−^ mice. Chronic pain mechanisms are known to emerge between Day 7 and Day 14 following surgery or inflammation in rodents (64, 65), and pain is generally considered as chronic beyond Day 14. (66). To model this, we injected Complete Freund’s Adjuvant (CFA) into the hind paw and monitored the animals over a 28-day period, assessing both thermal and mechanical sensitivity. The study included 20 WT mice (10 males, 10 females) and 19 THIK2^−/−^ mice (10 males, 9 females). To support the validity of the chronic inflammatory pain model, we confirmed that both genotypes developed thermal hyperalgesia by Day 22, consistent with the onset of central sensitization typically observed after more than 14 days of inflammation in mice. This supports the validity of the model studying chronic inflammatory. Notably, THIK2^−/−^ mice exhibited sustained thermal hyperalgesia throughout the testing period relative to baseline (compared to Pre-CFA, Day 0), with a significant difference between THIK2^−/−^ and WT mice emerging as early as Day 8 and lasting until the end of the test (Day 28) (Fig. 6A). In the Von Frey test, THIK2^−/−^ mice displayed sustained mechanical hyperalgesia throughout the 28-day period too (compared to Pre-CFA, Day 0), with significantly greater sensitivity compared to WT mice between Days 8 and 20. While WT mice progressively recovered and returned to mechanical baseline threshold by Day 28, THIK2^−/−^ mice failed to do so (Fig. 6B). A sex-specific difference was observed within the THIK2^−/−^ group at Day 2 post-CFA, with female mice displaying higher mechanical sensitivity than males (Supplementary Figure S4B). Overall, THIK2^−/−^ mice exhibited a more pronounced inflammatory hyperalgesia phenotype than control mice in response to thermal and mechanical stimuli, particularly during the period associated with the development of central sensitization (Day 8 to Day 16) (Fig. 6 A-B). Notably, thermal hyperalgesia persisted beyond Day 20 in THIK2^−/−^ mice, indicating a transition to a chronic pain state (< Day 20) (Fig. 6A). Importantly, the increased pain sensitivity observed in THIK2^−/−^ mice is not due to enhanced peripheral inflammation as paw thickness measured regularly over the 28-day following CFA injection was found to be thicker only on the day following injection and then similar throughout the test between genotypes (Fig. 6C).

**Fig. 6.**
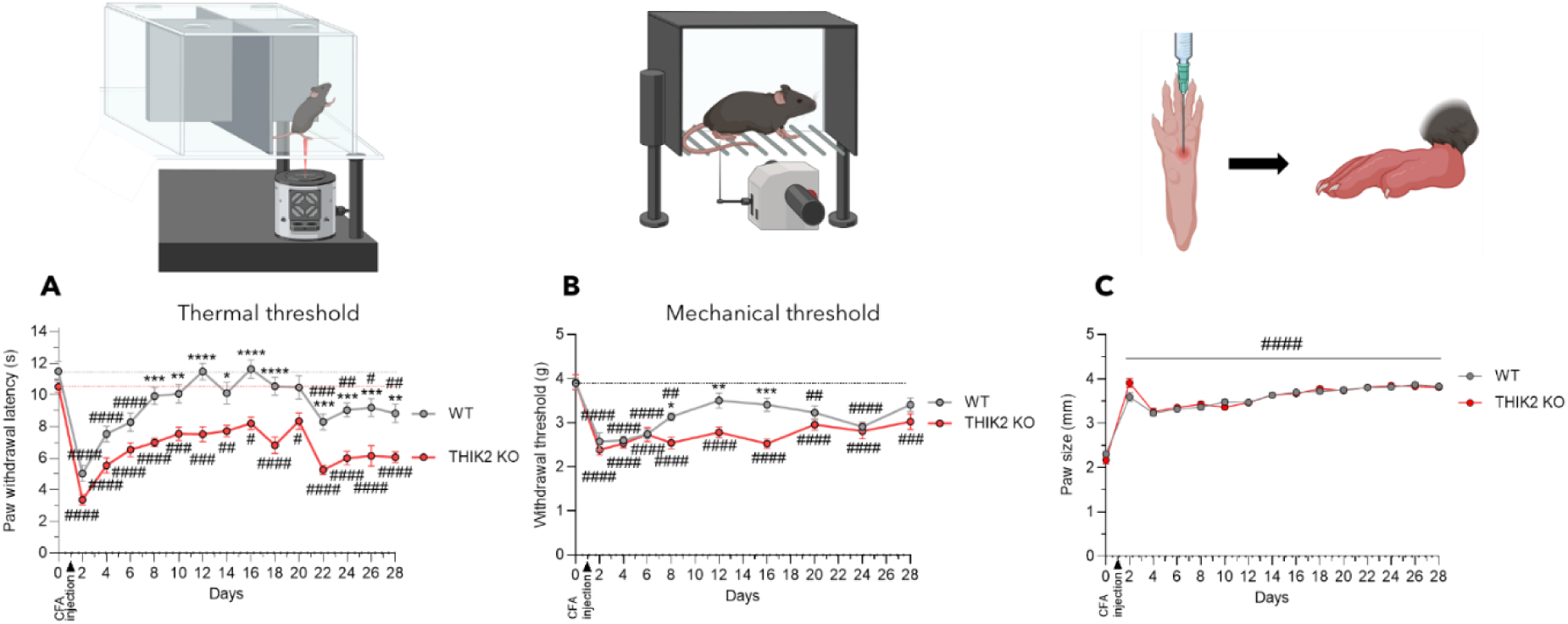
Chronic inflammatory pain of WT and THIK2^−/−^ in response to CFA injection. **A-B.** Evolution of paw withdrawal latency during 28 days of WT (n=20) and THIK2^−/−^ mice (n=19) in response to thermal stimuli (50 % power) in Hargreaves test (A) or mechanical stimuli (1g-7.5g) in dynamic Von Frey test (B). **C.** Thickness evolution of the hind pawbefore and after Complet Freund’s Adjuvant paw injection. **\*** means compared to THIK2^−/−^, **#** means compared to J0. Mean ± SEM, Two-way ANOVA with Bonferroni’s multiple comparison test, *p<0.05. **p<0.01. ***p<0.005 and ****p<0.001 and Two-way ANOVA with Dunett’s multiple comparison test to J0. #p<0.05. ##p<0.01. ###p<0.005. ####p<0.001.

## 4. Discussion

Chronic pain, particularly of inflammatory origin, remains a major clinical and public health challenge due to its high prevalence. Despite advances in therapeutics, current treatments often suffer from a lack of specificity, limited efficacy, or significant side effects, especially in the case of opioid-based therapies. In this context, the identification of new therapeutic targets beyond the opioid system has emerged as a promising direction in pain research. Alternative targets such as non-opioid membrane receptors including cannabinoid, glutamate, and neuropeptide receptors (e.g., substance P, NK receptors) as well as pro- and anti-nociceptive ion channels, are actively being explored and, in some cases, already applied in clinical settings (e.g., suzetrigine, a Nav1.8 channel blocker) (24). Additionally, cytokine receptors and immune-related mediators are under intense investigation, with several candidate molecules currently in development for the treatment of inflammatory pain (e.g., anti-TNF, anti-NGF, TAFA4) (67–69). In this landscape, our research investigates a completely novel and previously unexplored pathway in the field of nociception, with the objective of identifying alternative therapeutic strategies. Specifically, we examine for the first time the potential anti-nociceptive function of the potassium channel THIK2. which has not been studied to date in the context of pain modulation.

### THIK2 channel functions as an inhibitor of nociceptive transmission in C-fibers

Our results demonstrate that THIK2 and THIK1 are selectively expressed in a specific subpopulation of dorsal root ganglion (DRG) sensory neurons: the non-peptidergic C-fiber neurons (Figs. 1 & 2). These neurons are well recognized for their role in nociceptive detection and transmission, particularly during inflammatory pain (62). This precise expression pattern strongly suggests that THIK channels are directly involved in the early modulation of peripheral pain signaling. Electrophysiological recordings suggest that THIK2 modulates membrane excitability in sensory neurons (Fig. 3). The absence of major differences in resting membrane potential, input resistance, or single action potential (AP) properties indicates that THIK2 is not essential for passive membrane properties or the initiation of isolated APs. However, the increased number of APs in THIK2^−/−^ neurons during sustained stimulation highlights a tonic regulatory role of THIK2, likely acting as a brake to prevent pathological hyperexcitability in IB4+ non-peptidergic neurons. Other K₂_P_ family members have also been shown to act as critical brakes on neuronal excitability. For example, TREK1 and TRAAK significantly reduce the firing of peptidergic C-fiber nociceptors, thereby dampening pain signaling (32, 34). Similarly, both TREK2 and TRESK make substantial contributions to the resting membrane potential of dorsal root ganglion neurons, stabilizing their excitability under physiological and pathological conditions (31, 39, 42, 70, 71, 49).

These electrophysiological observations align closely with the behavioral hyperalgesia seen in THIK2 KO animals, supporting the idea that THIK2 normally helps maintain lower excitability thresholds in nociceptors. Indeed, THIK2^−/−^ mice display basal thermal hyperalgesia even without inflammatory stimuli, along with heightened spontaneous pain in models of non-chronic chemically induced inflammation (Fig. 4 & 5). Under chronic inflammatory conditions, these mice display marked mechanical and thermal hyperalgesia (Fig. 6), supporting a key role for THIK2 in polymodal nociceptors, particularly non-peptidergic C-fiber neurons, which are known to mediate slow-conducting pain and respond to diverse stimuli, especially in inflammatory states.

Further evidence for THIK2’s selective action in this neuronal subset comes from the observation that mechanical hyperalgesia in THIK2^−/−^ mice only manifests after 8 days of inflammation (Fig. 6B). Given that peripheral sensitization in mice begins around Day 2 (72), when normally innocuous Aβ fibers start to transmit pain signals, this delayed mechanical hyperalgesia suggests it is not due to Aβ fiber sensitization. Instead, it likely arises from increased excitability of polymodal, slowly conducting C-fibers (non-peptidergic). Together, these findings support the hypothesis that THIK2 functions as a protective modulator, reducing neuronal hyperexcitability in polymodal C-fiber nociceptors under both naïve and inflammatory conditions.

### Advantages of targeting THIK2 over other channels for pain modulation

These findings align with the already known role of K_2P_ in pain physiopathology, through their role in regulating resting membrane potential and controlling neuronal excitability. Disruption of these potassium conductance, whether through genetic deletion, transcriptional regulation, or pharmacological inhibition, can promote neuronal hyperexcitability leading to acute and chronic pain.

Genetic ablation of TRESK enhances mechanical and cold sensitivity, whereas sensitivity to heat remains largely unaffected (41, 45). Transcriptomic analyses revealed that TRESK exhibits preferential expression in NP1 and NP2 subtypes of sensory neurons that are implicated in itch signal transmission. Llimos-Aubach *et al.,* has shown in 2025 that TRESK deletion also exacerbates chronic itch in mouse models of allergic contact dermatitis, dry skin, and imiquimod-induced psoriasiform dermatitis, resulting in a significantly increased scratching behavior that develops earlier and is more robust. TRESK has a minimal expression in NP3 neurons and likely plays a minor role in their excitability (49). Although THIK2 is also expressed in NPEPs neurons, like TRESK, it does not appear to modulate itch in the same way as TRESK. However, several subpopulations of NPEP neurons exist form NP1 to NP3 (73). It would therefore be interesting to determine whether TRESK and THIK2 are expressed in the same subpopulations or in distinct ones. Furthermore, it would be useful to know the relative expression levels of these two channels in the DRG versus the TG, to explore whether, for example, TRESK, known to be involved in migraine, is more implicated in facial sensitivity, whereas THIK2, more highly expressed in the DRG, might be responsible for sensitivity in the rest of the body.

TRAAK is ubiquitously expressed across all sensory fiber types (31, 33). Genetic ablation of TRAAK leads to heightened thermal and mechanical pain sensitivity, demonstrating its tonic inhibitory role in nociception (34). TREK1 and TREK2 are both present in C-fiber nociceptors (40), with TREK1 preferentially localized to peptidergic (PEP) (74). Both channels contribute critically to the electrical stability of dorsal root ganglion neurons, as their blockade increases neuronal excitability and produces hyperalgesia in rats (75). Specifically, TREK1 modulates heat-evoked responses in C-fibers and TREK1 KO mice display pronounced heat hypersensitivity (32). While TREK2 KO animals exhibit both thermal and mechanical hypersensitivity (35). Several small-molecule activators of TREK and TRAAK channels have been evaluated for their analgesic potential, yet most candidates have been limited by unwanted systemic effects. Although compounds such as RNE28, LPS2336 effectively enhance K_2P_ channel currents and reduce nociceptor excitability, they concurrently induce hypotension, sedation, and muscle weakness in preclinical models (38, 76). These off-target physiological consequences underscore the difficulty of selectively modulating broadly expressed K_2P_ channels and suggest that shifting focus toward a more peripherally restricted target such as THIK2 may prove a more fruitful strategy for developing effective, side-effect–sparing analgesics.

Recent high-resolution transcriptomic (58) and proteomic (77) studies in humans have confirmed the expression of THIK2 in human DRG, providing a critical translational link. Remarkably, THIK2 is expressed in the same nociceptor subtypes in humans as in mice, particularly in nociceptive neurons. Furthermore, it ranks among the top ten most abundantly expressed ion channels in human DRG neurons (58, 77, 78). This conserved expression pattern across species strengthens the relevance of our murine findings and highlights THIK2 as a key candidate in human pain physiopathology. Given the behavioral data obtained in mice, THIK2 emerges as a promising target for the development of new treatments for chronic pain in humans.

RNAscope analyses demonstrated strong co-expression of THIK1 and THIK2 in P2RX3+ neurons, suggesting potential formation of functional heterodimers, as previously proposed (61). The slight downregulation of THIK1 expression in THIK2^−/−^ mice indicates a limited compensatory mechanism, reinforcing the validity of this KO model to study THIK2-specific roles without major confounding effects. To dissect the potential synergistic roles of THIK1 and THIK2 in sensory neurons, we propose generating a THIK1-THIK2 double-knockout mouse model. By comparing this double KO to each single KO, we can assess compensatory changes in the expression of other K_2P_ channels (e.g., TREK1. TREK2. TRESK) at the mRNA levels. Concurrently, a comprehensive battery of pain assays (thermal, mechanical, and inflammatory tests) will reveal how loss of both THIK isoforms alters nociceptive thresholds and behavioral hypersensitivity. This approach will clarify whether THIK1 and THIK2 function redundantly, additively, or synergistically to maintain neuronal excitability and modulate pain. This is important to consider, as THIK1 has been implicated in microglial surveillance and activation of the NLRP3 inflammasome in the central nervous system (79, 80). In this context, inhibiting THIK1 rather than activating it is recommended to reduce inflammation. Therefore, activating THIK1 to relieve pain could be counterproductive if it simultaneously triggers central inflammation that negates the analgesic benefit.

### Futures directions

Overall, our findings reveal a novel and specific role for THIK2 in the regulation of thermal and inflammatory nociception, particularly through non-peptidergic C-fiber neurons. The minimal compensatory changes in other ion channel expression observed in THIK2-/- mice support a non-redundant function for THIK2, positioning it as a promising therapeutic target, especially in the context of chronic inflammatory pain and opioid-sparing strategies.

Several future directions emerge from this study. Identifying selective modulators of THIK2, validating them in diverse preclinical pain models, and evaluating their safety and efficacy in clinically relevant conditions will be crucial steps. While the structure of the THIK1 channel has recently been resolved by cryo-EM, that of THIK2 remains undetermined. However, the high degree of similarity between the two channels—apart from differences in their trafficking— suggests that structurally conserved regions could be targeted for the development of channel activators (81–84). Moreover, a deeper understanding of the molecular mechanisms regulating THIK2 expression and function in sensory neurons will be essential to optimize future therapeutic strategies. Given the expression of THIK2 in various tissues beyond the peripheral nervous system, including several brain regions and the kidney (85–87), it would be worthwhile to explore its potential roles in other physiological processes. Interestingly, THIK2 KO mice do not display major phenotypic abnormalities, suggesting the existence of compensatory mechanisms in other system than sensory system or that THIK2’s role may be context-dependent. Sex-disaggregated analysis reveals that female THIK2^−/−^ mice show greater thermal hypersensitivity at 50% intensity, while both sexes are affected at 60 %. These findings, can reflect possible interactions between THIK2 function and hormonal or sex-specific signaling in nociceptive neurons.

To better define the specific contribution of THIK2 within NPEP neurons, a conditional KO approach targeting this particular subpopulation would be highly informative. This could involve crossing floxed THIK2 mice with a Cre-driver line specific to NPEPs such as Mrgprd-Cre (available on Jax.org), or alternatively, using viral vectors such as rAAV2/6 to deliver a dominant-negative form of THIK2 directly into IB4-positive neurons (88). Such strategies would help clarify whether THIK2 deletion in NPEPs alone is sufficient to reproduce the pain-related phenotypes observed in global knockouts. From a pharmacological standpoint, this model would also be critical in assessing whether peripheral inhibition of THIK2, without crossing the blood-brain barrier, is sufficient for effective pain relief. In parallel and in the absence of a selective pharmacological activator for THIK2, a knock-in (KI) approach either constitutive or conditional, could offer valuable insight. This is especially relevant given that our previous studies have identified mutations capable of rendering THIK2 either non-functional (61) or gain-of-function (89, 90).

In conclusion, our study explores the foundations of a new-fields of research in nociception, focusing on a little-known potassium channel. It highlights the importance of exploring not only pro-nociceptive mechanisms but also the endogenous negative regulators of pain, such as THIK2. This approach has the potential to fundamentally reshape chronic pain management, by addressing its functional causes rather than just its symptoms.

## Acknowledgements

Authors thank Drs J. Noël, A. Baron, M. Meynier, M. Toft, A. Moqrich, M. Poet and C. Sanchez for comments and discussions, J. Fassy for help in molecular biology, N. Guy and P. Pozzo Di Borgo for animal facility, C. Laredo for secretarial assistance and Dr Y-L. Shen for itch studies. We also thank the microscopy facility from the « Institut de Pharmacologie Moléculaire et Cellulaire » part of the « Microscopie Imagerie Cytométrie Azur» GIS IBiSA labeled platform and SABLES Platfomes financed by the European Union through the European Regional Development Fund ».

This work was supported by FRM, LabEx ICST, CNRS, INSERM and ANR K2PAIN.

## Author contributions

N.G. designed, performed, and analyzed molecular, behavioral and electrophysiological experiments. F.L., D.B. and F.C.C helped in experimental designing. D.B. and N.G. wrote the manuscript. S.F., F.L. C.C.C and F.C.C. participated in the critical reading of the manuscript. F.C.C. helped in patch-clamp experiments. T.L. and Y-L.S. helped in behavioral experiments. M-E.K. helped in genotyping and S.G. helped in molecular experiments, dissection and DRG culture. F.F. participated in the creation of the THIK2 KO mouse line.

Authors declare no conflicts of interest.

