## Supplementary figures 1 to 6 for "Two pore domain THIK2 channel is involved in acute and chronic pain signal regulation"

### Supplementary Informations

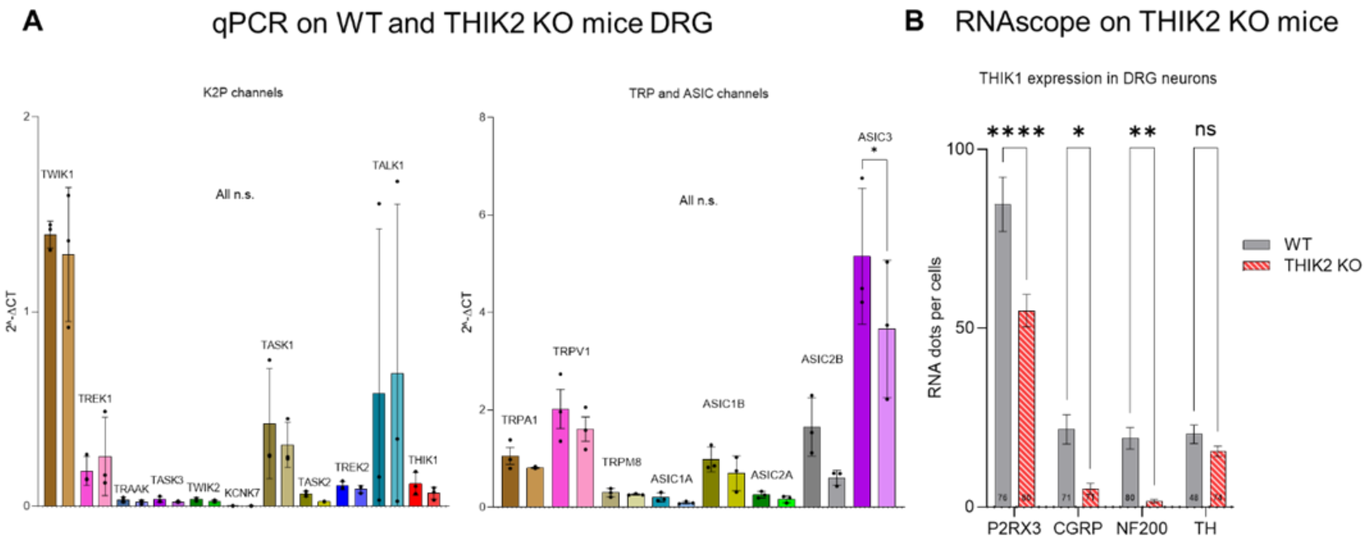


**Figure S1. Comparison of ion channels mRNA expression levels in mouse DRGs.**

**A.** Biomark qPCR on WT (n=3) and THIK2 KO (n=3) mice DRG showing expression levels of $K_{2P}$, ASIC and TRP channels mRNA in total DRG. The results are represented in $2^{-\Delta CT}$ and are normalized by the expression of the TOP1 reporter gene. Mean ± SEM, one-way ANOVA with with Sidak's multiple comparisons test, *p<0.05. **p<0.01. **B.** Quantification of THIK1-encoding RNA spots by RNAscope analysis in DRGs from 3 WT mice and 3 THIK2 KO mice. Mean ± SEM, two-way ANOVA with with Sidak's multiple comparisons test, *p<0.05, **p<0.01 and ****p<0.001. n represent the number of counted neurons; for P2RX3 n=76 WT, n=80 THIK2$^{-/-}$; for CGRP n=71 WT, n=71 THIK2$^{-/-}$; for NF200 n=80 WT, n=86 THIK2$^{-/-}$; for TH n=48 WT and n=74 THIK2$^{-/-}$.

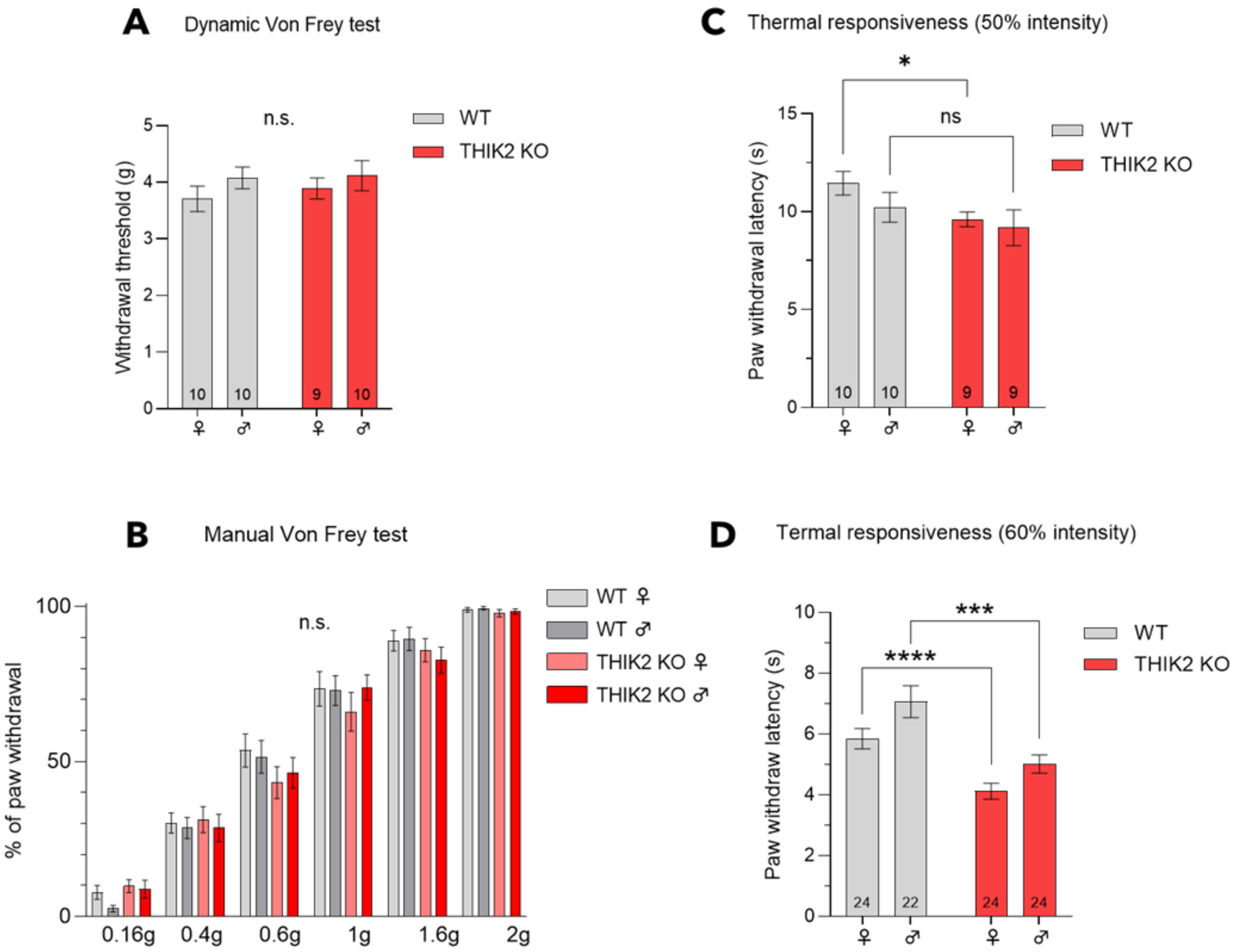


**Figure S2. Acute mechanical and thermal sensitivity of WT and THIK2$^{-/-}$ Mice with gender distinction. A-B.** Paw withdrawal threshold of male and female THIK2$^{-/-}$ mice compared to male and female WT mice in response to mechanical stimuli in dynamic von Frey test (1-7.5g) (A) or manual Von Frey test (0.16-2g) (B) (mean ± SEM, Unpaired T-test and two-way ANOVA). **C-D.** Paw withdrawal latency of THIK2$^{-/-}$ mice compared to WT mice in response to thermal stimuli, 50% (C) or 60% (D) intensity in Hargreaves test (mean ± SEM, Multiple T-test, *p<0.05. ***p<0.005. ****p<0.001). Different N are shown on graphs.

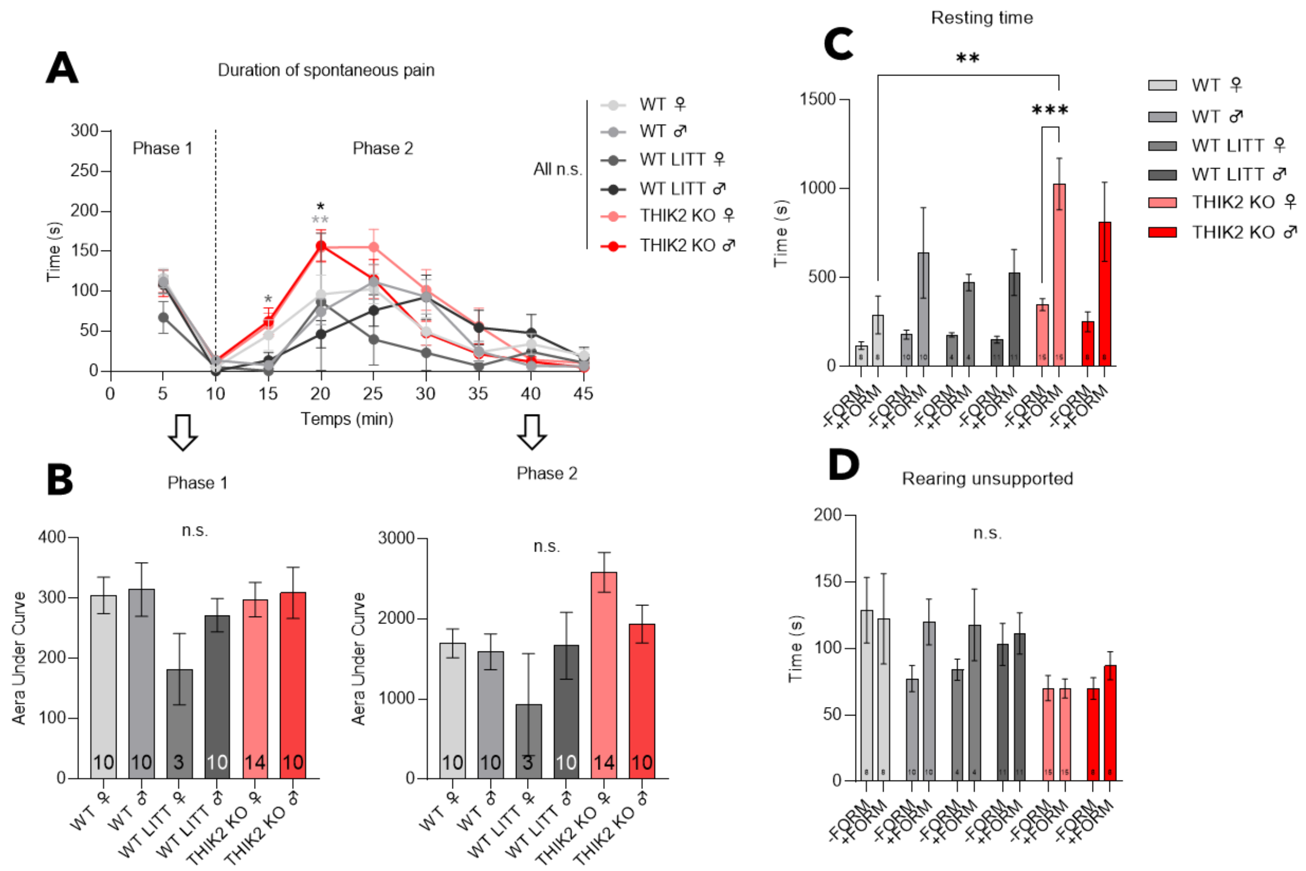


**Figure S3. Acute inflammatory spontaneous pain of WT, WT Littermates and THIK2$^{-/-}$ Mice with gender distinction.**

**A.** The pain behaviors measured in the formalin spontaneous pain test were licking, biting, shaking and paw lifting. The test lasted 45 minutes after the injection of 15µL of formalin into the left hind paw (Mean ± SEM, two-way ANOVA with with Tukey's multiple comparisons test, *p<0.05. **p<0.01). **B.** Area under curve form behaviors were separated into two phases: Acute phase between 0-10min and inflammatory phase 10-45min (Mean ± SEM, Multiple t-test). **C-D.** Post-formalin recovery was tested by placing the animals in a Phenotyper® observation cage 15min after the end of the formalin test, resting/prostration time and time spent rearing were measured (Mean ± SEM, two-way ANOVA with with Tukey's multiple comparisons test, **p<0.01 and ***p<0.005). Different N are shown on graphs.

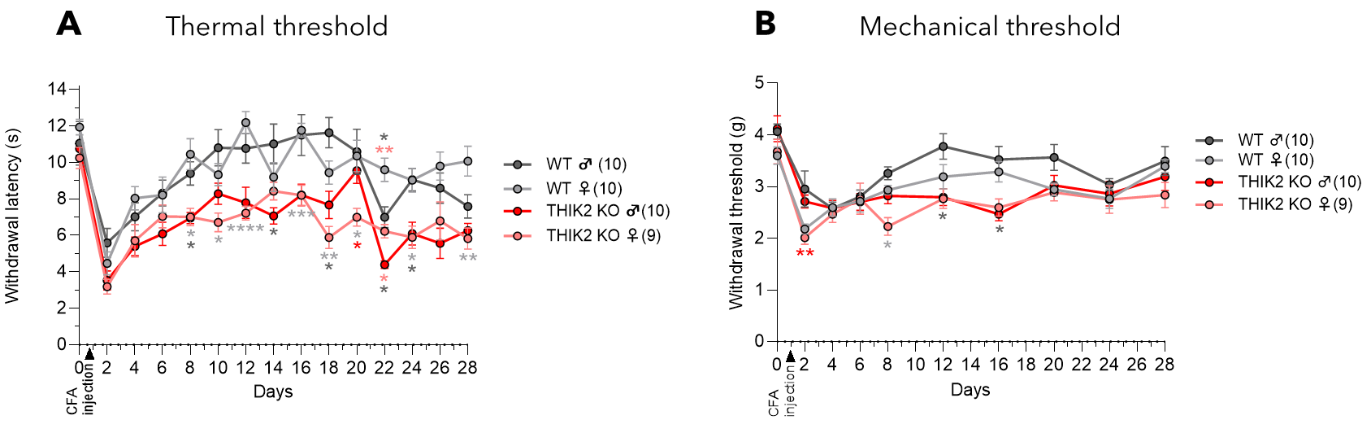


**Figure S4. Chronic inflammatory pain of WT and THIK2$^{-/-}$ in response to CFA injection with gender distinction.**
**A-B.** Evolution of paw withdrawal latency during 28 days of male and female WT and THIK2$^{-/-}$ mice in response to thermal stimuli (50 % power) in Hargreaves test (A) or mechanical stimuli (1g-7.5g) in dynamic Von Frey test (B) before and after Complet Freund's Adjuvant paw injection; * means Compared to THIK2-/-, # means Compared to J0 (mean ± SEM, Two-way ANOVA with Bonferroni's multiple comparison test, *p<0.05. **p<0.01, ***p<0.005 and ****p<0.001 and Two-way ANOVA with Dunett's multiple comparison test to J0. #p<0.05, ##p<0.01, ###p<0.005, ####p<0.001). N=10 males WT, N=10 females WT, N=10 males THIK2$^{-/-}$, N=9 females THIK2$^{-/-}$.

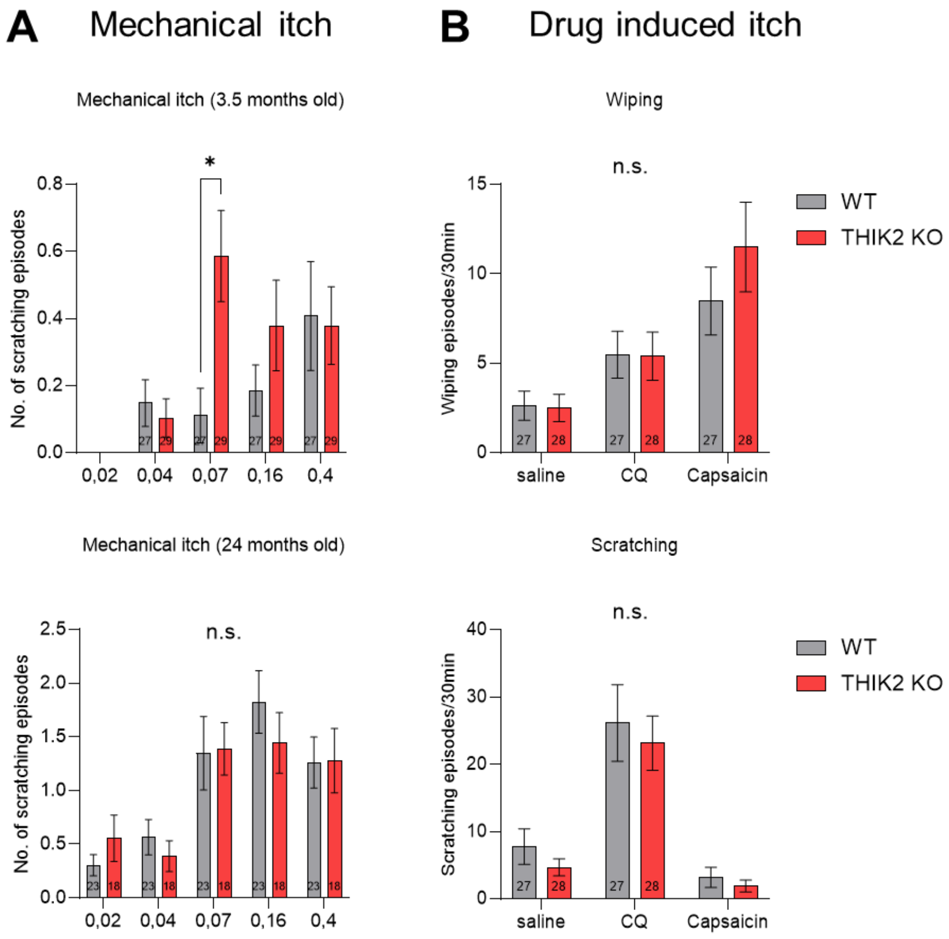


**Figure S5. Mechanical and drug induced itch in WT and THIK2$^{-/-}$ Mice.**
**A.** Number of scratching episodes after mechanical itch induced by Von Frey hairs (0.02-0.4g) at 3.5 months and 24 months (Mean ± SEM, Multiple paired t-test, *p<0.01). At 3,5 months old: N=27 WT and N=29 THIK2$^{-/-}$ mice. At 24 months old: N=23 WT and N=18 THIK2$^{-/-}$ mice. **B.** Number of wiping and scratching after saline, cloroquine (CQ) and capsaicin injection (Mean ± SEM, Multiple paired t-test). N=27 WT and N=28 THIK2$^{-/-}$ mice.

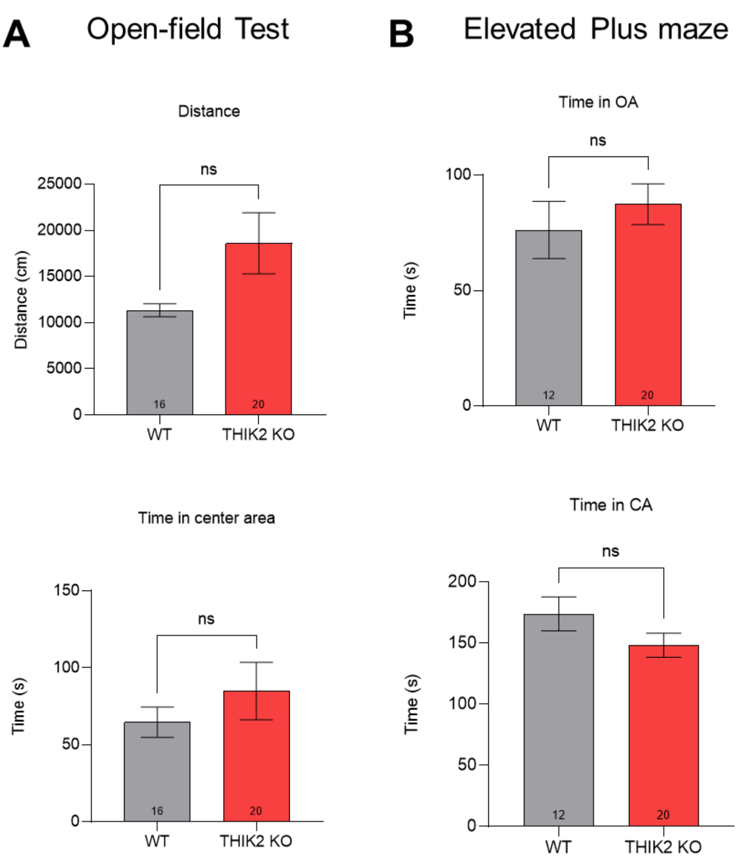


**Figure S6. Anxiety behavior and locomotor activity of WT and THIK2$^{-/-}$ Mice.**

**A.** Distance and time spent in center area by THIK2$^{-/-}$ and WT mice in open field test (Mean ± SEM, Unpaired t- test). N=16 WT mice and N=20 THIK2$^{-/-}$ **B.** Time spent in open arms (OA) and closed arms (CA) by THIK2-/- and WT mice (Mean ± SEM, Unpaired t- test). N=12 WT mice and N=20 THIK2$^{-/-}$ mice.
